# Scaffold Protein RACK1 inhibitor compounds prevent the Focal Adhesion Kinase mediated breast cancer cell migration and invasion potential

**DOI:** 10.1101/2021.06.13.448244

**Authors:** Hemayet Ullah, Nagib Ahsan, Sivanesan Dakshanamurthy

## Abstract

Scaffold protein RACK1 mediates cancer cell migration mostly through regulation of focal adhesion (FA) assembly by promoting a focal adhesion kinase (FAK) activation downstream of the integrin clustering and adhesion at the extracellular matrix (ECM). Here we demonstrated the efficacy of our recently developed RACK1 Y246 phosphorylation inhibitor compounds (SD29 and SD29-14) to inhibit the migration and invasion of MCF7 and MDA-MB-231 breast cancer cell lines. Using multiple assays, our results confirmed that inhibitor compounds effectively prevent the filopodia/lamellipodia development and inhibits the migration of breast cancer cells. A mechanistic model of the inhibitor compounds has been developed. Migration and invasion capabilities of the cancer cells define the metastasis of cancer. Thus, our results suggest a potential therapeutic mechanism of the inhibitors to prevent metastasis in diverse cancers.

## INTRODUCTION

Breast cancer is the most commonly diagnosed cancer in females and the leading cause of cancer death worldwide [1]. Though significant advances have been made in breast cancer treatment and therapies, still the major impediments in the advancement stem from the inability to counter the metastasis process that attacks distant organs. It is established that the migration and invasion of cancer cells to distant organs, not the localized cancer cells, is the major cause of cancer deaths. The majority of patients (>90%) succumb to cancer due to systemic cell metastases [2]. At present, there is no effective drugs or countermeasures available to target these specific invasive cells. Effective countermeasures to prevent the migration/invasion of cancer cells is sought after to limit the spread of cancer to distant organs.

Receptor of activated protein C kinase 1 (RACK1) is a highly conserved intracellular adaptor protein that plays a prominent role in many cancer cells including breast cancer cell invasion and migration that usually lead to the metastasis process [3–6]. RACK1 protein has been found to regulate the scaffolding of signaling proteins at the receptors. This regulation is particularly important in dynamic processes such as cell migration, cell adhesion and cell spreading through establishing contact with the extracellular matrix and growth factor receptors at the adhesion sites [7]. These processes include several signaling pathways, and require significant and well-orchestrated cross-talk between cell surface receptors and elements of the cell cytoskeleton [8].

RACK1 has been reported to regulate cancer cell migration and invasion through integration of three different cellular pathways- a physical interaction with Src kinase to modulate this key regulator of cancer cell migration[9]; through interaction with the Focal Adhesion Kinase (FAK) to regulate the polarity and direction-sensing of cancer cells from apical basal to front-rear direction [10]; and finally, by promoting the Epithelial to Mesenchymal transition (EMT) - a process known to promote the migration and invasion of cancer cell [11]. Src is a well-known regulator of cell adhesion, cell spreading and cell migration [12]. Src activity is inhibited by the binding of RACK1 but loss of RACK1 can prevent the transport of Src to specific cellular compartments where Src can function [13]. RACK1 is also found to regulate the assembly and functioning of the Focal Adhesions which are large dynamic macromolecular assemblies with both mechanical components and cell signaling components in the cancer cell migration process [10, 14]. Suppression of RACK1 expression disrupts FAK activity, cell adhesion and cell spreading [15]. EMT is an important process during cancer metastasis when epithelial cells lose the apical-basal polarity and cell-cell adhesion and transform the cell into invasive mesenchymal cells. RACK1 has been implicated in the EMT and in the invasion process in diverse cancer cells like in the esophageal squamous cell carcinoma [16]; in human glioma [11]; in prostate cancers [17]; in breast cancers [18–21]; in colon cancer [22]; in cervical cancer [23] and in non-small cell lung cancer [24]. An overarching mechanism of RACK1 to mediate so many diverse cancer types can be summed up by the dominant role RACK1 plays in the migration and invasion of cancer cells (reviewed in [4]).

As RACK1 has been established as the key regulator of cancer cell migration, invasion that are prerequisite for the metastasis process, it is expected that development of any RACK1 protein functional inhibitor compounds will be the key to prevent cancer cell metastasis. Based on our lab developed crystal structure of RACK1 protein, we have developed a class of small compounds targeting the functionally and experimentally confirmed key tyrosine phosphorylation site [25, 26]. Previously it was reported that upon activation by PKC, RACK1 colocalizes with Src at the plasma membrane where Src kinase phosphorylate RACK1 at the Y246 residue leading to the cell growth [27, 28]. Mutagenesis of the RACK1 on the same tyrosine residue resulted in the loss of interaction with other proteins, including with Src kinase [28–30]. As the compounds isolated to bind to the same tyrosine residue, it was shown previously that the treatment of the cells with the compounds could inhibit the signal induced same tyrosine residue phosphorylation which effectively inhibited cellular function of RACK1 [26].

As RACK1 has been established to regulate the cancer cell migration, adhesion, invasion, here we report that the application of RACK1 functional inhibitor compounds to two different breast cancer cell line. The results indicate that the treatment has effectively inhibited many of the hallmarks of migration/invasion of these cells as well as affecting the FAK expression and the EMT markers. It is hypothesized that within the cancer environment, RACK1 may act as a dynamic hub of signaling proteins regulating cell migration/invasion/metastasis.

## MATERIALS AND METHODS

### Cell Cultures, Reagents, and Antibodies

The MCF-7 and MDA-MB-231 triple-negative breast cancer (TNBC) cells were cultured in Dulbecco’s Modified Eagle’s medium (DMEM) media supplemented with 10% Fetal Bovine serum (FBS), penicillin (100 IU/ml)-streptomycin (100 μg/ml) and amphotericin B (2.5 μg/ml). Cells were seeded in 12-well plates or on 25 ml vented flasks, incubated in a humidified 5% CO2 incubator at 37°C. As indicated in the figure legends, for investigating invasion, cells were treated with Transforming Growth Factor (TGF-b) (Peprotech, Cranbury, NJ) at indicated concentrations (2ng-10ng/ml). The molecular structure; M.W.; chemical characteristics of the RACK1 inhibitor drugs are detailed in (Ullah et al., 2019). The drugs were dissolved in DMSO and were used from 1 to 50 uM concentration. All reagents were from Sigma-Aldrich (St Louis, MO) or Fisher Chemicals (Fairlawn, NJ) unless otherwise indicated. The mouse monoclonal RACK1 antibody raised against amino acids 131-317 of RACK1 of human origin was purchased from Santa Cruz Biotechnology (Cat# sc-17754, 1:2000 dilution). phospho-FAK (Tyr397) rabbit polyclonal antibody (dilution 1:1000; # 700255, Life Technologies Corporation (Thermo Fisher). α-Tubulin (11H10) Rabbit mAb (cat# 2125S), Cell Signaling. Mouse TRITC secondary antibody (cat# A16016) - ThermoFisher. Rabbit FITC secondary antibody (cat# T-2765)-ThermoFisher; Antifade with DAPI (cat# P36966)-ThermoFisher. Cell staining was done with Giemsa stain (1:20 diluted in water) or with 0.5% crystal violet dissolved in 25% Methanol. Corning Matrigel Basement Membrane Matrix (cat# 356234) was used diluted 1:6 with serum free media.

### Protein extraction, Immunoblots

Cells were washed three times with cold PBS. Lysates were isolated in lysis buffer (Cell Signaling, Danvers, MA) supplemented with protease inhibitor (Sigma-Aldrich), phosphatase inhibitor cocktail A and B (Santa Cruz Biotechnology, Dallas, TX), and N-Ethylamaleimide (Sigma-Aldrich, St. Louis, MO). The concentration of all proteins was measured using Bio-Rad’s Bradford dye following the manufacturer’s instructions. After 50 microgram of lysates with 1X loading dye (biorad) supplemented with 50 μl/ml βME (beta-mercaptoethanol) were boiled for 10 minutes, chilled lysates were loaded on the BioRad’s 4-12% precast Bis-Tris polyacrylamide gel, Proteins were then transferred onto a precut PVDF membrane (Biorad) using Biorad electro-transfer set up. The membrane was then blocked with 5% non-fat milk in PBST (1X PBS with 0.1% Tween 20) for one hour, three washes were done for 10 minutes each by using PBST (1X PBS with 0.1% Tween 20). Washed membrane was incubated overnight at 40Cwith the indicated diluted primary antibody, followed by incubation with a horseradish peroxidase-coupled secondary antibody. After washing three times with PBST, the protein expression was visualized by Biorad’s Clarity ECL reagent using Protein Simple FluorChem M system.

### Immunofluorescence microscopy

Immunofluorescence experiments were performed as previously described (Ullah et al., 2019). Briefly, the cells on glass coverslips in 12 well plate were incubated with indicated treatments or with DMSO for 96 hours. Then cells were washed 3X with PBS, fixed with 4% PFA-PBS for 15 minutes. The permeabilization was done at room temperature with 0.5% Triton X-100 in PBS (pH 7.4) for 10 min. Cells were washed 3X with ice cold PBS. For blocking, the cells were incubated with 1% BSA, 22.52 mg/ml glycine in PBST (PBS with 0.1% Tween 20) for 30 min at room temperature. The washed cells were incubated in a humified 4°C chamber overnight with the indicated antibody (1:500-both together for two protein detection) in PBST with 1% BSA. The cells were washed three times in PBS, 5 min each wash and then Incubated cells with the anti-rabbit FITC and anti-mouse TRITC conjugated secondary antibody (1:1000) in 1% BSA for 1 h at room temperature in the dark. After washing three times with PBST for 5 minutes each, the cells were mounted on slide using Antifade mounting media with DAPI (4-6-diamidino-2-phenylindole; ThermoFisher). After sealing the coverslips with clear nail polish, the slides were left at 4°C overnight for curing. Imaging was performed using an Eclipse Ti 2000 laser-scanning confocal microscope (Nikon CSU series Spinning Disk confocal microscope) with FITC, TRITC, and DAPI filter sets.

### Migration and invasion analysis

Cells were grown to almost confluency on a 6 well plate, in 1% serum DMEM media supplemented with 4 ng/ml of TGF-β1 for the duration of the experiment. The wound was initiated by displacing a line of monolayer cells with tip of a sterile one ml pipette tip. The displaced line was photographed at time zero and allowed to incubate in the same media with DMSO or with SD29-14 (10 μM) for 48h. The same area was then photographed to assess the migration of cells towards the displaced area.

For assessing directional invasion of cancer cells, modified Boyden dual chamber assay was done. Coring 6.5 mm Transwell with 8.0 um pore polycarbonate membrane insert in 12 well plates were used (Sigma Aldrich, cat# CLS3422). The insert membrane was treated for optimal cell attachment. A total of 1.0 × 105 cells were cultured in serum free media with DMSO or with SD29 (1 μM) for 24 hours in the upper chamber cell inserts. The lower chamber contained 10% serum media as chemoattractant. After 24 hours of growth inside the CO2 incubator, the cells and media from the upper chamber is removed and wiped with a cotton swab. The cells attached to the lower side of the membrane is washed 3X in PBS and fixed with 4% PFA. Then the membrane was stained with 0.5% crystal violet in 25% methanol. The cells stuck to the membrane were photograph with a 10X objective of a microscope.

### LC-MS/MS analysis and database search

SDS-PAGE gel slices from each sample were subjected to in-gel trypsin digestion according to the established protocol. Tryptic peptides were further purified by Pierce™ Graphite Spin Columns following manufacturer’s instructions. The dried tryptic peptides were resuspended in water with 0.1% formic acid (v/v) and equal volume of tryptic peptides was subjected for LC-MS/MS analysis. The LC-MS/MS was performed by using an LTQ Orbitrap XL mass spectrometer (Thermo Fisher Scientific) coupled to a Prominence Nano LC (Shimadzu Corporation, Columbia, MD, USA) using the Xcalibur version 2.7.0 (Thermo Scientific). The detail MS/MS analysis condition, instrument setup, and LC condition were described in the earlier published protocol (Lin et al., 2017; PMID: 29100346).

Peptide spectrum matching of MS/MS spectra of each file was searched against the UniProt human database (TaxonID: 9606, downloaded on 10/25/2017) using the Sequest algorithm within Proteome Discoverer v 2.4 software (Thermo Fisher Scientific, San Jose, CA). The Sequest database search was performed with the following parameters: trypsin enzyme cleavage specificity, 2 possible missed cleavages, 10 ppm mass tolerance for precursor ions, 0.02 Da mass tolerance for fragment ions. Search parameters permitted dynamic modification of methionine oxidation (+15.9949 Da) and static modification of carbamidomethylation (+57.0215 Da) on cysteine. Peptide assignments from the database search were filtered down to a 1% FDR. The relative label-free quantitative and comparative among the samples were performed using the Minora algorithm and the adjoining bioinformatics tools of the Proteome Discoverer 2.4 software.

## RESULTS

Various specialized membrane protrusions are the hallmark of cancer cells that facilitate cell to cell communications and dissemination of metastatic cells. There are many different types of cell protrusions like Invadopodia, lamellipodia, filopodia, podosomes are defined in the cancer cells [2, 31]. In order to investigate whether the RACK1 inhibitor compounds have any role in the development of the cellular membrane protrusions in the breast cancer cells, MCF-7 cells were treated with the compounds (Fig.1).

**Figure 1.**
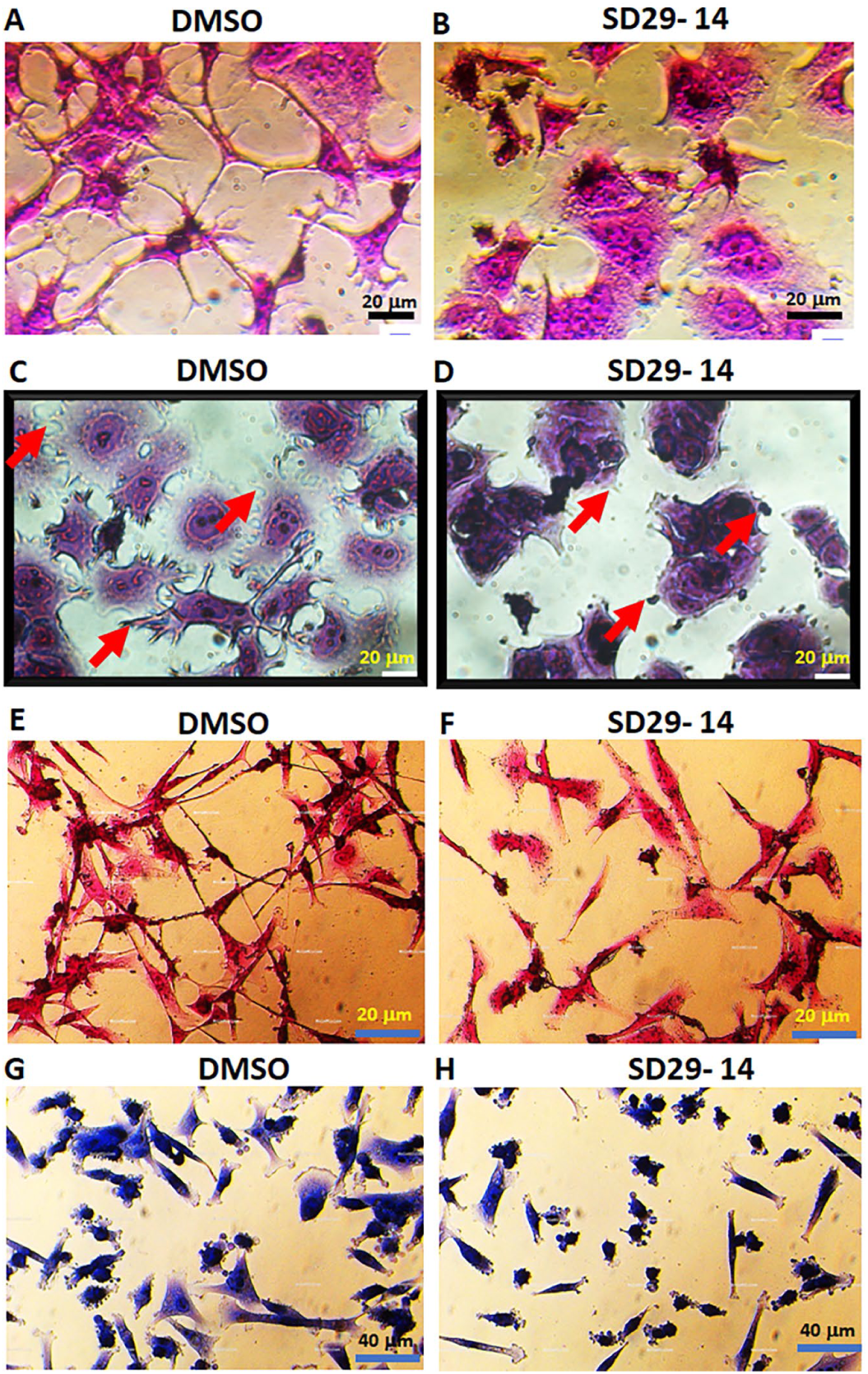
Functional Inhibitor compound of RACK1 prevents the development of membrane protrusive structures in MCF-7 and MDA-MB-231 cells. DIC photomicrograph of MCF-7 cells grown for 24h (no serum) followed by 48h growth with DMSO **(A)** or with 1 μM SD29-14 **(B)** in the presence of 10 ng/ml TGF-b1. Cells were fixed with 4% formaldehyde and stained with crystal violet (0.5 %) in 25% methanol. The cells were photographed with 40X objective. Scale bar = 20 μm. **(C) & (D):** MCF-7 cells treated with DMSO **(C)** or SD29-14 **(D)** for 48h were then allowed to grow on no serum basement matrix matrigels (1:6 diluted with serum free media) for 20h. After removing media, matrigels, PBS washed cells were stained with Giemsa stain (1:20 diluted in water) and were photographed under a 40X objective. Scale bar = 20 μm. **(E) & (F):** MDA-MB-231 cells were grown for 72h in the presence of DMSO **(E)** and 50 μM SD29-14 **(F)** and then PBS washed and fixed followed by stained with crystal violet (0.5 % in methanol). Cells were photographed with a 40X objective. Scale bar = 20 μm. **(G) & (H):** MDA-MB-231 cells were grown for 16h with DMSO **(G)** or with 1 μM SD29-14 (H), PBS washed, fixed 4% formaldehyde and then stained with Giemsa stain. Cells were photographed under 20X objective. Scale bar = 40 μm

The cells were treated with SD29-14 (1 □M) for 72h in the presence of Transforming Growth Factor (TGF-beta1-10 ng/ml) to facilitate cell proliferation and differentiation. As can be seen from Fig 1, the DMSO treated control cells (Fig. 1-panel A) developed extensive network like cell membrane protrusions while cells treated with the inhibitor compound SD29-14 (Fig. 1-panel B) almost no or rudimentary type protrusions are seen on the cell membranes

We also used MCF-7 cells grown on basement membrane matrix- matrigel and 5 ng/ml TGF-b1 so that the cells can attach, proliferate and develop membrane protrusions to allow migration of the cells in the matrix. As can be seen in Fig. 1, control cells treated with DMSO (panel C) that were grown on Matrigel developed extensive lamellipodia with short filopodia like extensions at the leading edge of the cells (red arrows). However, treatment of the cells with the SD29-14 almost completely inhibited and sometimes appeared as abortive (red arrows) appendages from the cell membrane. No clear lamellipodia like protrusions were seen in the inhibitor treated cells. Though in this experiment a higher concentration of inhibitor (100 uM) was used to allow effective diffusion of the compound inside the Matrigel, similar inhibitions of the cellular protrusions were also seen from low concentration of the compound (data not shown- used in the patent application).

The similar set of experiments were done with the triple negative breast cancer cell line MDA-MB-231 which is known for their aggressive growth and proliferation. As can be seen in Fig. 1, treatment with DMSO (panel E) resulted in the development of extensive cell to cell connection and membrane protrusions indicating their potential for invasive capabilities. Whereas, treatment with SD29-14 (50 uM) did not develop such membrane protrusions but rather developed diminished type of lamellipodia like protrusions (panel F). Also, when the MD-231 cells were grown for a shorter time in the presence of TGFb1 (10 ng/ml), the DMSO treated cells developed numerous conspicuous podosomes, lamellipodia like protrusions (panel G) while SD29-14 (1 uM) treatment developed reduced podosomes and rarely any lamellipodia. These indicate that without the inhibitor compound, the cancer cells rapidly acquired the competence for migration which was diminished in the presence of the inhibitor compound.

Collectively, the results clearly show that without the drug treatment, the breast cancer cells developed extensive membrane protrusions that have led to development of filipodia and lamellipodia like structures making the cell competent for invasion and migration. Whereas the drug SD29-14 treated cells show an inhibition of such membrane protrusions and as such were not able to develop the filipodia and lamellipodia like structures. The RACK1 functional inhibitor compound-based prevention of migration/invasion competent cells supports the previous findings where RACK1 upregulations have been implicated in diverse cancer cells migration and invasion [3, 4].

During tumorigenesis, the epithelial-mesenchymal transition (EMT) plays a crucial role in migration and invasion of various cancers. EMT plays an essential role in tumor invasiveness and metastasis in cancer progression and the process has been shown to be regulated by the RACK1 protein [11, 16]. The motile mesenchymal cells can move (migrate) into surrounding tissue, even to remote tissues. Therefore, assays were done to investigate the migration and invasion potential of MB-231 cells with or without the SD29-14 treatment. A wound was created by sliding a tip of a 1 ml plastic pipette tip on the surface of almost 100 percent confluent cells and the size of the wound was measured at 0h which were identical for both the DMSO and the SD29-14 (10 □M) treated cells. However, after 48h of growth in the incubator, the DMSO treated cells migrated to significantly cover up the area of clearing (Fig. 2A) but in the presence of the SD29-14 (Fig. 2B), the cells failed to migrate to cover the clear area created by the wound indicating that the presence of SD29-14 may have suppressed the cells migration potential.

**Figure 2.**
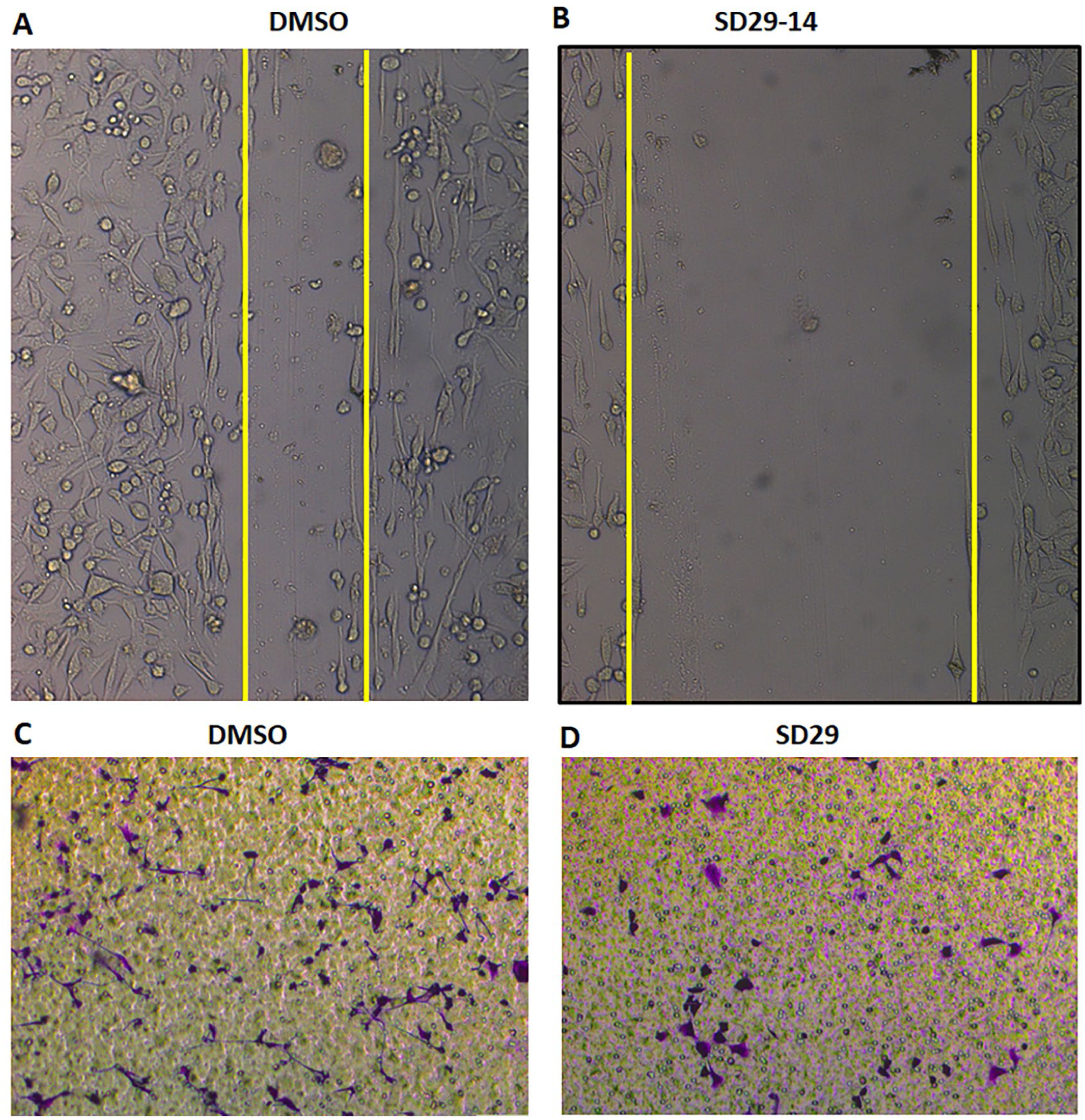
Treatment with RACK1 functional inhibitor compound to MDA-MB-231 cells decreased cell motility. **(A)** and **(B):** Phase-contrast micrographs of living cultures of TNBC MDA-MB-231 cells in a wound-healing assays performed in the presence of DMSO **(A)** or with SD29-14 **(B)** 10 μM. The cells were treated with 4 ng/ml of TGF-β1 for the duration of the experiment. Almost confluent cells cultured in media with 1% serum and the wound was initiated by making a line displacing the cells with an 1ml pipette tip. Cells were photographed at 0h (not shown) and at 48h with a 10X objective. **(C) and (D):** Inhibitory effects of the compound on transwell migration and invasion abilities of MB-231. **(C)** The effect of inhibitor compound-SD29 (1 μM) on MDA-MB-231 cells was investigated by transwell migration assay without Matrigel in the presence of DMSO (C) or 1 μM of the inhibitor compound SD29 (parent compound for the analog SD29-14). Cells were allowed to grow with the compound on the upper chamber for 24h and migration towards the bottom chamber with serum was assayed by staining the chamber membrane with crystal violet (0.5% in 25% methanol). The migrated cells stuck on the lower side of the membrane were photographed (10X objective) and representative images are shown in the panel **C** and **D**.

The transwell invasion (Boyden Chamber) assay was used to test the invasive potential of the MB-231 cells in response to migration inducers. During this assay, cells were placed on the upper layer of a cell culture insert with a permeable membrane (8 um pore size). When the MB-231 cells (0.5 × 10 ^5) were placed on the upper chamber with serum free media containing TGFb1 and no SD29 for about 24 hours, a large number of cells were captured on the bottom side of the membrane as the cells tried to move towards the lower chamber with serum (Fig. 2C). The invasive action of the cells towards the media with serum was visualized by crystal violet stain. Under similar conditions, very few cells were observed on the membrane when SD29 was added to the cells on the upper chamber (Fig. 2D). Collectively, these findings indicated that both SD29 and SD29-14 can be used to have a significant inhibitory role in the regulation of invasive breast cancer. Note that both drugs were found to inhibit RACK1 Y248 phosphorylation [26].

### SD29-14 blocks Tyr^397^ phosphorylation of FAK in MCF-7 cells

It was shown in the MCF-7 cells that FAK phosphorylation is a pre-requisite for cell adhesion and interaction of RACK1 with FAK was required for the adhesion and subsequent migration of the cancer cells [15]. Based on earlier results, it was expected that inhibition or disruption, if not complete blockage, of FAK phosphorylation and improper localization of FAK could in turn inhibit metastasis. This may happen as FAK phosphorylation disruption would inhibit the proper development of lamellipodia and the cellular polarity that are associated with motility and invasiveness characteristic of cancer cell migration. Therefore, we set our experiments to test whether the RACK1 inhibitor compounds could prevent the Tyr397 phosphorylation of FAK in the MCF-7 cells. In this immunofluorescence experiment, The MCF-7 cells were grown for 96h in the presence of TGF-b1 (5 ng/ml). The cells were fixed and probed with two antibodies- anti-pY397-FAK and anti-RACK1. The pY397-FAK expression is ascertained by staining the primary antibody with a FITC tagged secondary antibody (green) and the RACK1 primary antibody is detected with TRITC tagged secondary antibody (Red). As can be seen from Fig 3, the treatment of the cells without the inhibitor compounds (left panel) clearly shows extensive expression of Y397 phosphorylation of FAK (green) at the nascent protrusive nascent adhesions that are needed for adhesion and migration, whereas RACK1 is seen to be expressed in the cytoplasm as well as in the protrusive structures. On the other hand, treatment with the inhibitor compound SD29-14 resulted in the significant inhibition in the expression of the pY397-FAK protein with concomitant reduction in the RACK1 expression (right panel). As discussed above, without the pY397-FAK present, the connection between the ECM (extra cellular matrix) and cell interiors will be disrupted which will inhibit the cell’s potential to migrate by initiating the cell’s adhesion to the ECM. The significant result indicates that the RACK1 inhibitor compound will be useful in the inhibition of pY397-FAK based cancer cell adhesion and migration.

**Figure 3.**
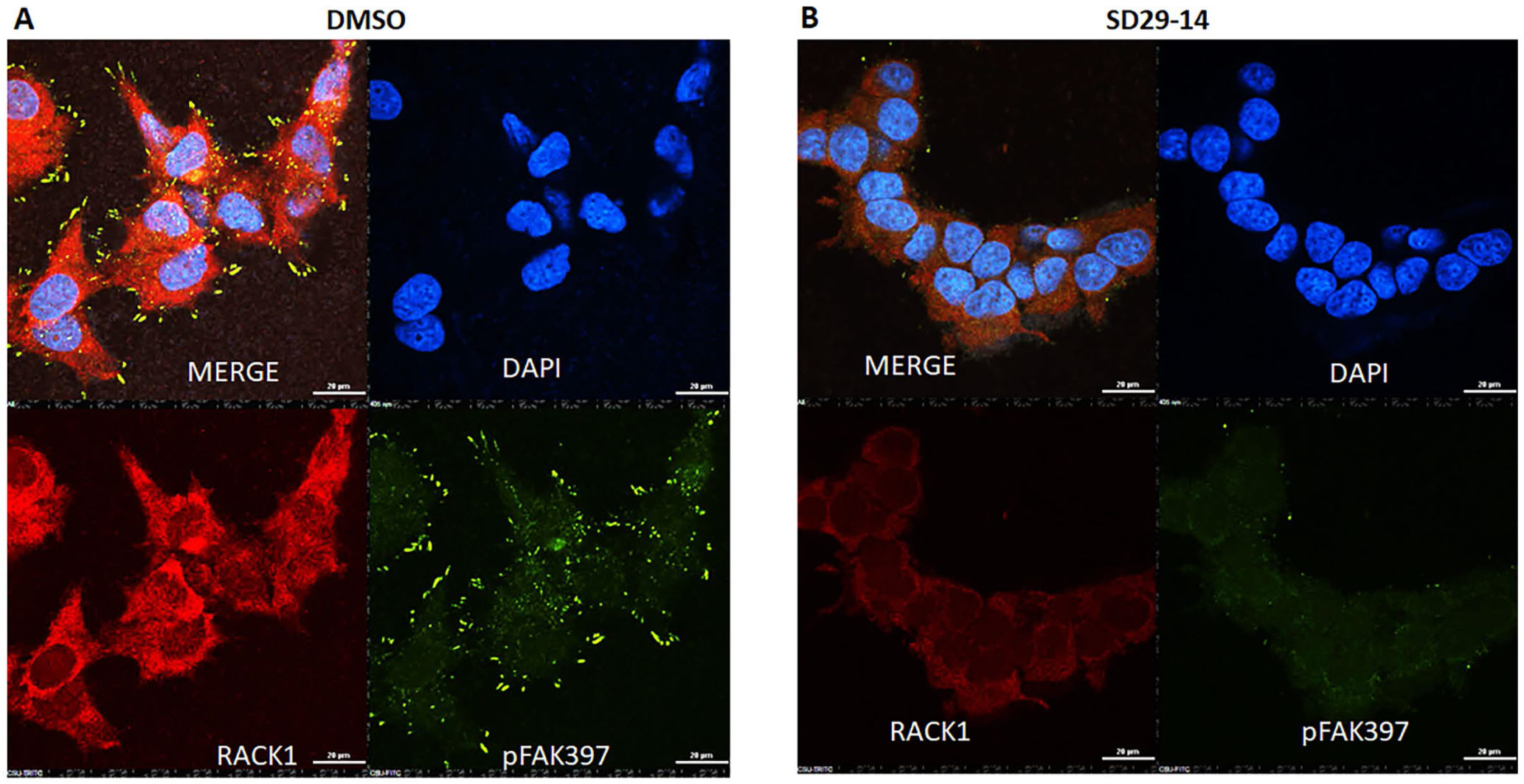
Immunofluorescence evidence for RACK1 inhibitor compound induced inhibition of FAK Y397 phosphorylation needed for migration in MCF-7 cells. The cells were treated with either DMSO **(A)** or with 50 μM inhibitor SD29-14 **(B)** for 96 hours. Fixed cells were stained with anti-pFAK397-FITC (green) and with anti-RACK1-TRITC (Red). The DAPI (blue) stain was used to stain the nuclear DNA. Fluorescence cells were observed under 60X objective. Scale bar 20 μM. Fluorescence images from at least 6 different spots on the slides were taken and a represented image is presented here.

### The Compound Inhibited RACK1 Expression

We also investigated the potential of the inhibitor compounds in the inhibition of RACK1 expression in both the MCF-7 and in the MB-231 cells. The cells were grown for 96 hours with different concentration of the inhibitor compounds and with the control DMSO. The lysates were probed with an anti-RACK1 antibody. As can be seen from the Fig. 4, the expression of the RACK1 in both the cancer cell lines was inhibited significantly with compound concentration of as little as 1 uM. This inhibition of RACK1 expression by the compound could make the cells not develop the protrusive nascent adhesions as seen in Fig.3 during the growth of the cells. This effect on the both the cancer cell lines indicate that the compounds can potentially be used on diverse cancers where upregulation of RACK1 protein has been found to induce migration of cancer cells.

**Figure 4.**
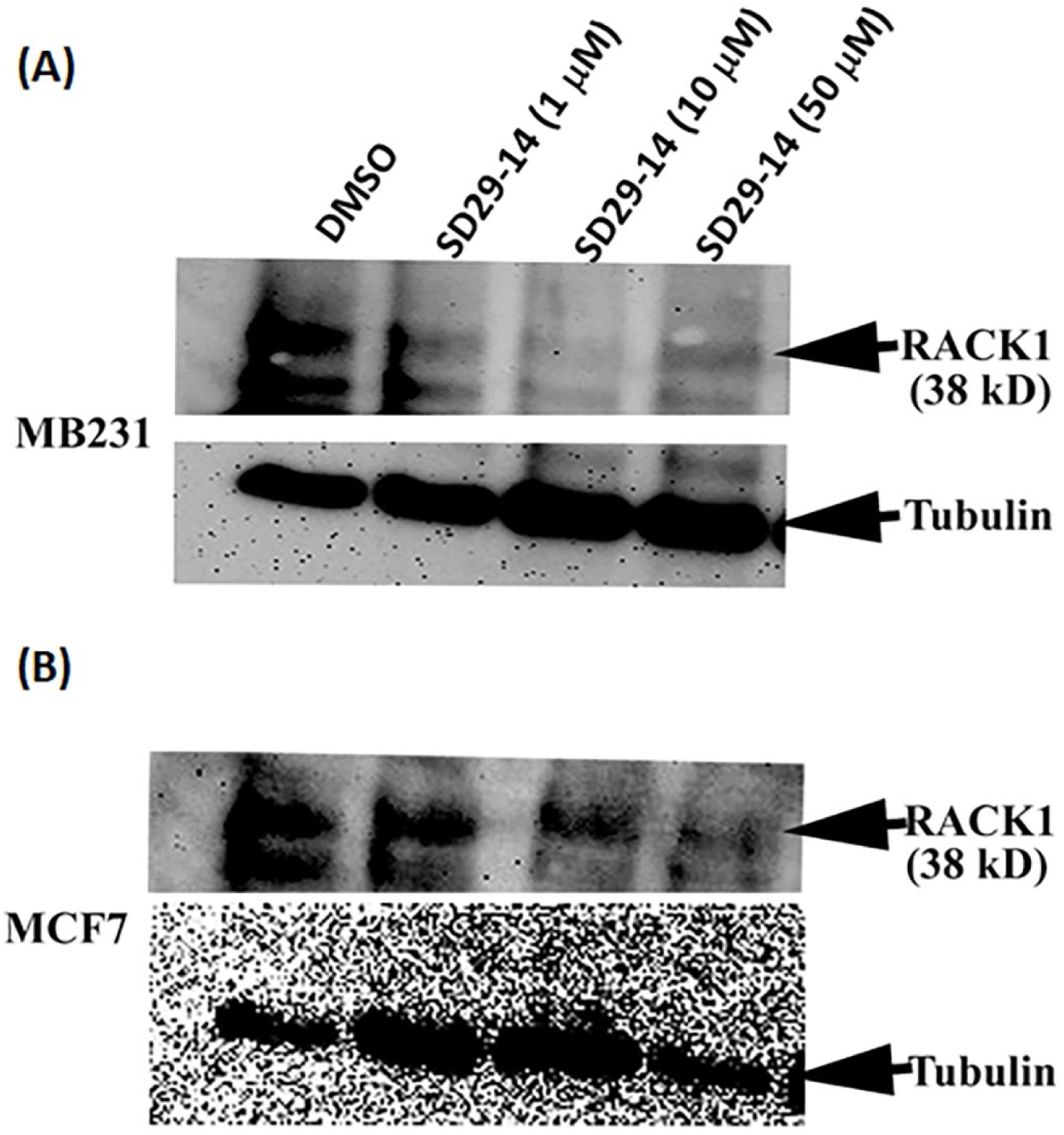
RACK1 functional inhibitor compound inhibits concentration dependent RACK1 protein expression in MB-231 (upper panel) and in MCF7 (lower panel) breast cancer cells. Different concentration of SD29-14 drug treatment (1, 10, and 50 mM) in cells grown for 96 hours. DMSO was used as drug solvent control. The lysates were probed with a mouse anti-RACK1 antibody. The anti-tubulin antibody was used as loading control on the same membrane after stripping off previous RACK1 antibody.

In order to unravel other proteins that may be involved in the RACK1 inhibitor compound induced process of inhibiting cancer cell migration, we used a proteomic analysis experiment where differentially expressed bands on SDS-PAGE gels were subjected to in-gel trypsin digestion according to the established protocol. A total of 211 peptides corresponding to 44 unique proteins were successfully identified and quantified by LC-MS/MS analysis from the SDS-PAGE gel samples (Supplementary Table 1). Every single peptide/protein identified with 1% FDR confidence level for the sequence assignments indicates very high confidence. A principle component analysis (PCA) (Figure 5A) of the total abundance of the proteins in each sample showed close clustering of the replication of each group, however distinct among the groups such as DMSO and SD29-14 treated samples. A heat map clustering analysis of the protein abundance among the samples further demonstrated the similar expression pattern of the proteins in each replication whereas distinct among the treatment (Figure 5B). Protein expression patterns of some target proteins such as ANXA1, ANXA2 VIM and RPS6 shows increased abundance in SD29-14 treatment whereas proteins like MB was not identified or shows decreased abundance in SD29-14 treated samples (Figure 5C). Annexin and vimentin are known interactor of RACK1, as such the inhibition of RACK1 may allow these proteins not to be sequestered by RACK1. Interaction of RACK1 with vimentin has been shown to regulate FAK mediated cancer cell migration [32]. Previously, it was found that RACK1 mediates tyrosine phosphorylation of Anxa2 by Src and promoted invasion and metastasis in drug resistant breast cancer cells [33].

**Figure 5.**
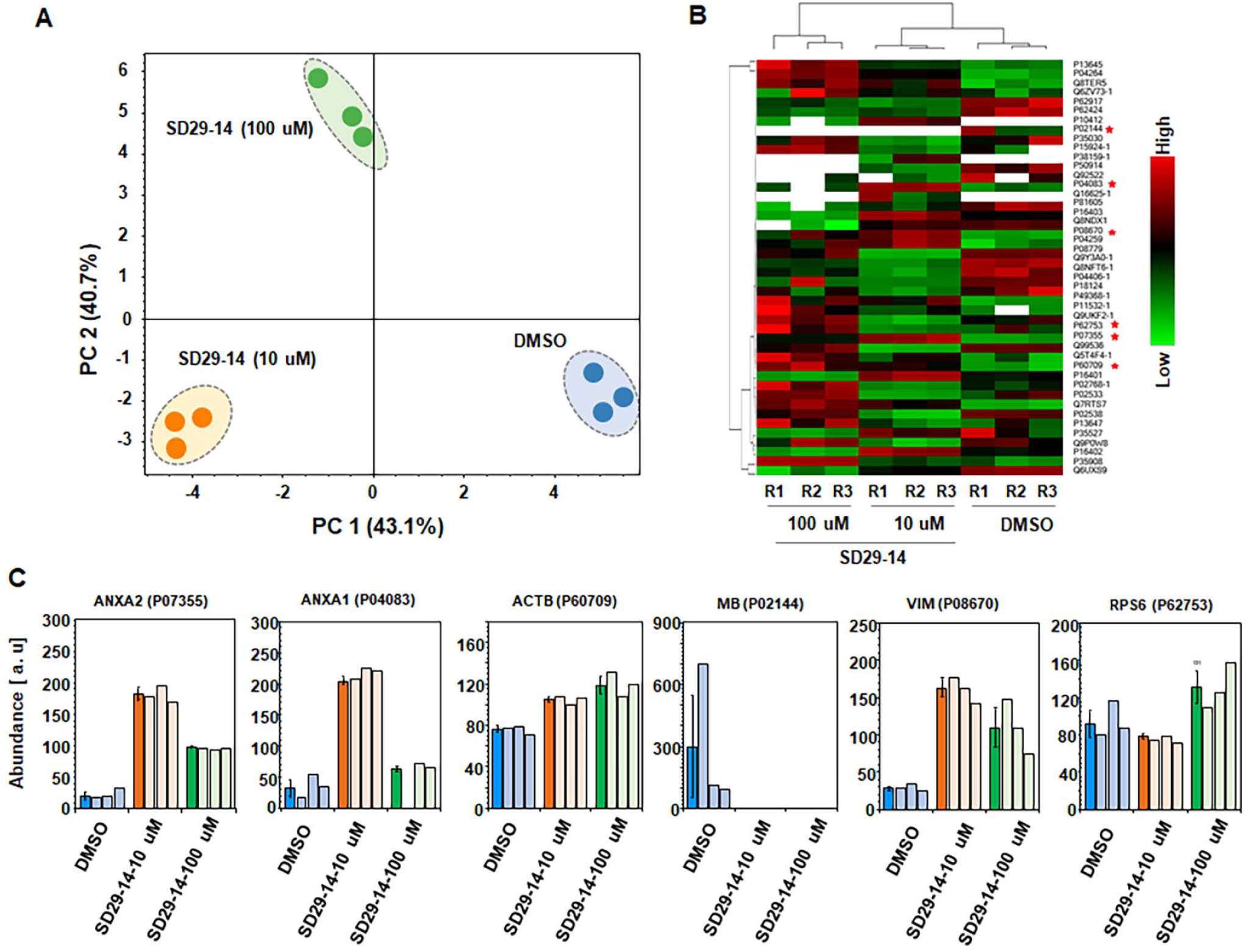
LC-MS/MS protein identification and quantification of DMSO, and SD29-14 treated samples. **A)** principal component analysis (PCA) of total protein abundance (peak area) data collected from each replication of DMSO, SD29-14 (10 uM) and SD29-14 (100 uM) treated samples. Data represents the close clustering of peptide peak areas from replicates of each condition and distinct among the three conditions. **B)** heat map analysis of the total 44 proteins identified and quantified from SDS-PAGE samples. Red star indicates the gene of interest. C, bar diagrams dark blue, orange, and green representing the average (n=3) protein expression peak area and light blue, orange, and green bars representing the peak are of each replication of the target proteins.

## DISCUSSION

Scaffold protein RACK1 is known to regulate diverse cancer cell migration mostly through regulation of focal adhesion (FA) assembly. This assembly mainly happens through the activation of focal adhesion kinase (FAK) downstream of integrin clustering and adhesion at the extracellular matrix (ECM). Here we report the application of our recently developed RACK1 inhibitor compounds [26] to two different breast cancer cells -MCF7 and MDA-MB-231 (TNBC) to inhibit adhesion, migration and invasion of the cells. Using assays to discern filopodia, lamellipodia and invadopodia development as well as invasion and migration assays coupled with immunofluorescence microscopy and mass spectrometry, here we show that the inhibitor compounds can effectively prevent the development of filopodia/lamellipodia and effectively inhibit migration of the breast cancer cells. RACK1 protein is known to use its WD repeats to mediate protein-protein interactions through recruiting signaling molecules to discrete cellular locations like at the cell membrane, ribosome, and nucleus [5]. RACK1 mediates cell spreading by establishing contact with the extracellular matrix (ECM) and ECM protein receptors-integrins at the focal adhesion sites [7, 8].

RACK1 is known be an indispensable component in a so-called direction sensing pathway that includes the integrin effector FAK (focal adhesion kinase) and PDE4D5 [3, 34]. FAK has been reported as a signaling switch for diverse cellular functions, including cell motility and directional control as well as tissue invasion [14]. When the Tyr397 residue of FAK is auto-phosphorylated (termed here as pFAK397), the development of a complex formation to assemble a focal adhesion structure or assembly has been described [4]. The complex has been associated with lamellipodia and other protrusive structures formation which are the characteristic of cancer cell migration and invasion (metastasis) process. It has been reported that phosphorylated FAK is associated with enhanced motility (migration potential) of several cancer types [35]. Autophosphorylation of FAK could be increased by RACK1 but when RACK1 expression was suppressed, FAK was not phosphorylated on Tyr-397 and was not responsive to stimulation by the IGF-I receptor in cells [15, 36]. Our results of RACK1 inhibitor compounds induced inhibition of pFAK397 expression could be the leading cause of the prevention of the migration of the cancer cells.

Involvement of the Src Family protein tyrosine kinases with the pFAK397 downstream of integrin clustering at the ECM can be the clue as for the mechanism of action for the inhibitor compounds. It is known that integrin clustering is sufficient to promote the phosphorylation of FAK (Focal Adhesion Kinase) on Tyr-397 that in turn recruits Src Family protein tyrosine kinases (Src-family PTKs) to the complex to regulate FAK activities [8]. However, phosphorylation of RACK1 on Tyr-246 is also required for the binding to the Src and binding of RACK1 to Src is essential to regulate FAK’s function [10, 12, 13]. Formation of the RACK1^Y246^-Src-FAK^397^ complex allows targeted phosphorylation of substrates at the focal adhesions and invadopodia by Src causing efficient invasion [8]. As the inhibitor compounds were developed targeting the same tyrosine phosphorylation site, abolition of RACK1 Y246 phosphorylation by the inhibitor compounds can potentially break the complex leading to the migration impairment. Previously it was convincingly showed that suppression of RACK1 led to the disruption in FAK activity, cell adhesion and cell spreading [10, 15, 36].

The inhibitor compound has been very effective in suppressing RACK1 expression. Indeed, altered RACK1 expression has been associated with the inhibition of diverse physiological functions that can lead to the development and maintenance of a variety of cancer hallmarks, including sustained proliferative signal, cell invasion and metastasis [6]. As discussed before, in addition to the breast cancer cells, RACK1 has been established as a regulator of diverse cancer cell proliferation, migration, invasion, and in metastasis [3, 4, 16, 37–40]. Elevated RACK1 expression in MCF-7 cells has been found to enhance the cell capacity for cell migration, growth, and invasion [5, 20], whereas down-regulation of RACK1 expression in breast cancer cells has led to the inhibition of proliferation, migration, and invasion *in vitro* and *in vivo* [19]. Therefore, RACK1 has been established as a potential drug target as a means to inhibit cancer cell migration and invasion. Here, we have shown that the drugs targeting RACK1 functional tyrosine phosphorylation can effectively act as an inhibitor of the migration and invasion in two phenotypically different breast cancer cells. Availability of the RACK1 inhibitor compounds will be instrumental in the elucidation of not only breast cancer cell migration, it will be equally useful in the diverse other cancer cells where RACK1 has been implicated in the effective migration, invasion, and eventual metastasis of cancer.

Structural studies has universally identified RACK1 as a ribosomal protein [41] whereas its interaction with diverse cellular signaling proteins [8] indicate that RACK1 can have both ribosome on and off function. In a recent review on RACK1 as a regulator of translation and breast cancer cell migration, Buoso et al. [6] presented a hypothesis explaining the protein’s role on and off the ribosome. It has been proposed that RACK1 may recruit the ribosome to focal adhesions (FA) for local translation of proteins to regulate cell spreading and migration [42]. However, Buoso et al. indicated that depending on mesenchymal and epithelial cancer cell profile, RACK1 may exert different functions in these two cell profiles. RACK1 may display a preferential ribosomal function in the epithelial profile, whereas holds a more structural role in the mesenchymal one, suggesting RACK1 extra-ribosomal functions correlate with a more aggressive breast cancer phenotype. Here the effect of the RACK1 inhibitor compounds on the migration and invasion was visibly effective on the more aggressive TNBC breast cancer line MB-231 which maintains a mesenchymal profile (Fig. 3). This indicate that the compounds may be more effective in inhibiting the extra-ribosomal function of RACK1; thereby, inhibit migration and invasion of more aggressive breast cancer cells.

Our proteomic studies of altered expression of Anxa2 along with Vimentin support the previous findings that RACK1 regulates migration/invasion of breast cancer cells by potentially acting as the signaling hub to bring various physiological regulators of migration together. In fact, Anxa2 protein phosphorylation by Src kinase only happens when RACK1 is present resulting in highly aggressive invasive breast cancer cells [33]. Using RACK1-Y246F mutation of RACK1 which is what the used SD29-14 inhibitor compound creates, the authors showed that rescued expression of WT RACK1, but not RACK1-Y246F mutant, in RACK1-silenced cells increased the number of metastases in the infected mice lungs compared to that of the control group [33]. Though Anxa2 was found to be expressed at high level with the inhibitor, it can be explained that the lack of interaction with RACK1 could potentially release the WT Anxa2 to a higher level without getting required phosphorylation by Src kinase to impart invasion potential to the cells. The study clearly indicated that RACK1 acts at a convergence point for mediating crosstalk among signaling molecules to regulate the development of the invasive/metastatic potential of breast cancer. In the same manner, it was found that expression of RACK1 is a prerequisite to form the intermolecular complex with Vimentin and FAK in the invading endothelial cultures in three dimensions [32]. Depletion of RACK1 decreased the association of vimentin and FAK, suggesting that RACK1 was required for stabilizing vimentin–FAK interactions during angiogenic sprouting [32]. The inhibitor compound SD29-14 most likely inhibited the intermolecular complex formation- releasing the Vimentin, without any required modifications like phosphorylation, that may ultimately resulted in the inhibition of invasive potential of the cells. Drugs that can interrupt RACK1 interaction with specific protein that is required for cancer cell invasion during metastasis can be very successful to inhibit directional invasion of cancer cells and this kind of directed interruption will be useful as it may inhibit only the specific function without affecting diverse other cellular cell signaling processes regulated by RACK1.

Based on the experimental results that included evaluations of the protrusive nascent adhesion sites and filopodia/lamellipodia development, migration potential, subcellular localization, and proteomic analyses, we have presented a potential model for the role of RACK1 in the process (Fig. 6). The model shows a nexus among Integrin, FAK, RACK1, and Src Kinase complex that extends to the cellular cytoskeleton to establish the polarity of the cancer cells. A detailed description of the model is provided in the legend of the Figure 6. It is of utmost importance to understand the complex process by which tumor cells invade and metastasize to distant sites utilizing multiple steps including local adhesion, migration, invasion and then dissemination of tumor cells through the lymphatic systems to the colonization of micro-metastatic lesions to ultimately into macro-metastasis.

**Figure 6.**
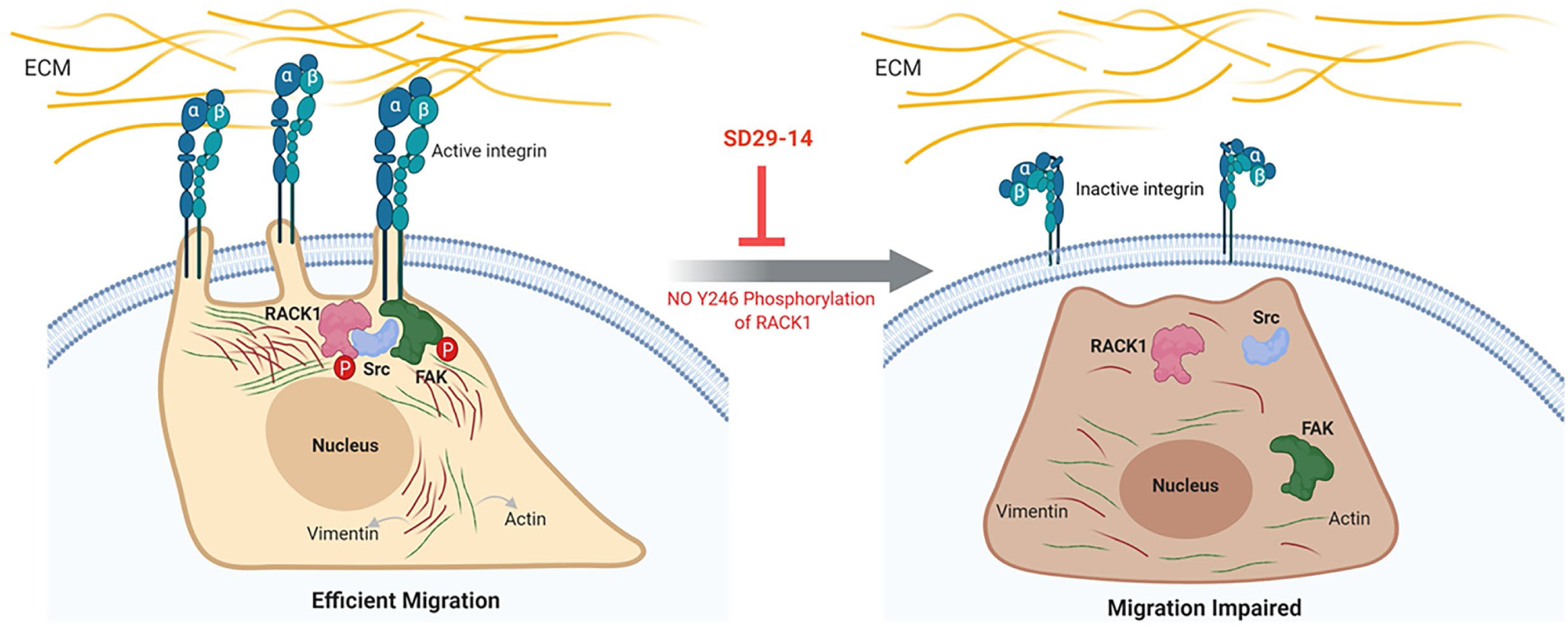
Proposed Model showing the role of the RACK1 inhibitor compounds on cancer cell invasion/migration. RACK1 acts as a versatile signaling hub for proteins regulating efficient invasion of cancer cells. RACK1 establishes contact with the extracellular matrix (ECM) and ECM protein receptors-Integrins at the Focal Adhesion (FA) sites. Integrin clustering is sufficient to promote the phosphorylation of FAK (Focal Adhesion Kinase) on Tyr-397. This in turn generates a binding site for the Src homology 2 (SH2) domain of Src Family protein tyrosine kinases (Src-family PTKs). The recruitment of Src-family PTKs to FAK is dependent on the initial phosphorylation of Tyr-397. Phosphorylation of RACK1 on Tyr-246 is required for the binding to the Src. RACK1 is also reported as the substrate of Src for this phosphorylation. Binding of RACK1 to Src is essential to regulate FAK’s function. Formation of phosphor-RACK1^Y246^-Src-FAK^397^ complex allows targeted phosphorylation of substrates at the focal adhesions and invadopodia by Src causing efficient invasion. Abolition of RACK1 Y246 phosphorylation breaks the complex and Src is free to phosphorylate substrate indiscriminately leading to migration impairment. In addition, interaction of RACK1 with vimentin is required to regulate FAK mediated cancer cell migration. Localization of RACK1 with vimentin in the perinuclear region provides potential for migration. For simplicity only few Actin (green thread) and Vimentin (brown thread) complexes in the migration process are depicted in the model. This model figure was created with BioRender.com.

## CONCLUSIONS

Development of any drug/compound that can inhibit any of the multiple steps will be a significant step in the fight against this key process that leads to almost 90% of the cancer-related mortality. Here we have shown that the functional inhibitor compound of the RACK1 protein that is known to be the key signaling hub for the cell migration and invasion regulators can be the significant development in this key cancer process. Finally, availability of the RACK1 inhibitor compounds will not only be beneficial for preventing the breast cancer cell migration and invasion, the compounds will also be useful in diverse other cancer cell migration/invasions where RACK1 has been implicated in the process. Future animal and clinical studies of the compounds will bring out the true potential of the compounds as a broad application drug for targeting diverse types of cancer cell migration and invasion.

## ACKNOWLEDGMENTS

The author Sivanesan Dakshanamurthy wishes to acknowledge the support by Georgetown University Medical Center - Lombardi Comprehensive Cancer Center, and CCSG grant P30 CA051008/CA/NCI NIH HHS/United States. We acknowledge Dr. Aniqa, Ph.D. at Harvard University for reading and editing the manuscript.

## SUPPLEMENTAL MATERIALS

**Supplementary Table 1.**

A proteomic analysis experiment to identify the other proteins that involved in the RACK1 inhibitor compound induced process of inhibiting cancer cell migration. Table shows the identified peptides corresponding to 44 unique proteins and quantified by LC-MS/MS analysis from the SDS-PAGE gel samples.

## Notes

### Competing Interest Statement

The authors have declared no competing interest.

### Summary of Updates

Figure 1 revised; Figure 4 revised.

